# An Ancient, Structurally Conserved Allorecognition Gene Complex Predates the Divergence of Major Cnidarian Lineages

**DOI:** 10.64898/2026.10.06.757116

**Authors:** Alberto M. Rivera, Jonah Petty, Andreas D. Baxevanis

## Abstract

The allorecognition system of the hydroid *Hydractinia symbiolongicarpus* plays an essential role in mediating self vs. non-self recognition among cnidarian colonies, analogous to the role of the major histocompatibility complex (MHC) in bilaterian immune systems. Both complexes have contiguous stretches of similar, highly duplicated genes, suggesting a homologous relationship. Protein structural prediction methods such as AlphaFold2 also indicate structural similarity between the proteins encoded within the *Hydractinia* ARC and the MHC. We have determined the structure of the ARC in the related hydrozoan *Podocoryna americana* to determine the degree of conservation of the cnidarian allorecognition complex. Protein structural prediction methods have revealed a number of candidate allorecognition proteins that form a long contiguous complex. Phylogenies suggest homology between cnidarian allorecognition genes, as well as syntenic conservation of this gene complex across hydrozoans. Ongoing comparative genomic analyses across bilaterian species aim to clarify the relationship between the ARC and the MHC, thereby informing our understanding of the evolutionary origins of bilaterian adaptive immunity.

## Introduction

Cnidarians are a key phylum for studying the evolution of novel tissues and functions in metazoans. This phylum, which includes jellyfish, anemones, corals, and hydroids, is the sister group to the bilaterians and, as such, its phylogenetic placement makes these animals useful models for understanding the evolution of essential functions in morphologically complex bilaterian animals. Cnidarians have distinctive life stages and life cycle strategies (Leclère et al., 2019), including free-living medusae and sessile colonies of polyps, within which individual polyps engage in specialized functions such as feeding and sexual reproduction (Cartwright et al., 2021).

An essential function of the colonial life stage is self/non-self recognition, or allorecognition (Nicotra et al., 2009). In colonial cnidarians such as *Hydractinia*, colonies typically grow on hermit crab shells, where space is limited and encounters with neighboring colonies are common. Such encounters may occur either between distinct colonies competing for space or within a single colony that has grown around the shell and contacts itself (Grosberg et al., 1996). The outcome of these interactions depends on the allorecognition response, where incompatible colonies are rejected through aggressive responses mediated by specialized stinging cells called cnidocytes. In contrast, contacts recognized as self result in fusion to maintain the integrity of the colony (Nicotra, 2019; Babonis, 2025). This allorecognition response is mediated at the molecular level by a family of highly polymorphic cell surface receptors called allorecognition proteins (ALRs).

Thirty-eight putative allorecognition proteins have been identified in *Hydractinia* and the genes encoding these proteins are clustered within a highly contiguous gene complex on a single chromosome called the allorecognition complex (ARC; Huene et al., 2022). This finding has prompted interest in the evolutionary relationship between the ARC and the major histocompatibility complex (MHC) of the bilaterian immune system. ARC-encoded ALRs share several structural features with bilaterian immune receptors (Huene et al., 2022), including one to four immunoglobulin-like domains that typically begin with a variable (V-set) Ig-like domain followed by intermediate (I-set) domains (Nicotra et al., 2009; Rosa et al., 2010). Many ALRs also possess a C-terminal fibronectin type III-like domain (FN3-like; Huene et al., 2022). ALRs and MHC proteins also share motifs associated with the immune response, including immunoreceptor tyrosine-based activation motifs (ITAMs) and immunoreceptor tyrosine-based inhibitory motifs (ITIMs) (Barrow & Trowsdale, 2006; Huene et al., 2022). In bilaterians, phosphorylated ITAMs initiate a downstream signal transduction pathway that activates an immune response, whereas ITIMs recruit inhibitory pathways that suppress these responses (Lanier, 2008).

These similarities extend beyond their overall protein domain architecture to the tertiary structure of the domains themselves. Ig-like domains in cnidarian ALRs show a high degree of tertiary structural conservation with bilaterian immunoreceptors despite having less than 20% sequence similarity (Huene et al., 2022), placing these comparisons within the “twilight zone” where sequence-based inference of homology becomes unreliable (Ponting, 2017). However, the observed conservation in protein domain structure and overall architecture suggests an evolutionary relationship between the cnidarian allorecognition system and the adaptive immune system of bilaterians. Across wide evolutionary distances, protein structure is often more highly conserved than amino acid sequence (Kresge et al., 2006), suggesting that methods based on protein structural similarity may be particularly informative (if not essential) for detecting distant homologous relationships of biological importance.

To date, the ARC has only been characterized in *Hydractinia*, limiting our understanding of how this gene complex originated and diversified across cnidarians. Analyzing the evolution of this essential gene family across multiple cnidarian taxa with distinct life cycles, colonial forms, and ecological strategies can clarify how allorecognition systems have evolved and how selection has acted on this gene family over time. Differences in the colonial life stage or the morphological organization of these colonies may impose distinct selective pressures on the allorecognition system and affect its evolutionary history. More broadly, analyzing the evolution of cnidarian allorecognition may shed light on the early origins of animal immune recognition which could, in turn, have potential relevance to immunology and human health. In this study, we have characterized the evolution of the ARC across multiple cnidarian taxa, establishing a framework for analyzing this gene complex in these close relatives of bilaterians.

## Methods

### Genome Selection and Protein Structure Prediction

We downloaded full-length chromosomal scale genomes for the following cnidarian species: *Podocoryna americana* (Chang et al., 2026), *Hydra vulgaris* (Kon-Nanjo et al., 2025), *Acropora millepora* (Fuller et al., 2020)*, Pocillopora damicornis* (Cunning et al., 2018), and *Rhopilema esculentum* (Nong et al., 2020; Goldfarb et al., 2025). These species were selected to provide representative sampling across major cnidarian clades, encompassing the hydrozoans, anthozoans, and scyphozoans.

To compare the evolutionary history of gene complexes and contiguous genomic regions, we limited our study to cnidarians with chromosomal-scale assemblies available in NCBI’s Reference Sequence Database (RefSeq; Goldfarb et al., 2025). We also used genome annotations and predicted protein sequences available from NCBI RefSeq in the course of our analyses. In cases where RefSeq annotations were not available, we instead used annotations provided by the group that generated the corresponding genome assembly. If no annotations were available, we performed *ab initio* prediction with BRAKER3 (Gabriel et al., 2024) to generate annotations. Since BRAKER3 annotations can be improved by incorporating a protein database, we used known metazoan proteins from OrthoDB (Kuznetsov et al., 2022), supplemented with sequences of published *Hydractinia* proteins (Huene et al., 2022; Schnitzler et al., 2024) to increase the likelihood of correctly annotating allorecognition-related genes given their high degree of divergence. We also prioritized RefSeq annotations to make the analysis as consistent as possible. The resulting predicted protein sequences were used for downstream homolog detection analyses.

Using AlphaFold2 (Jumper et al., 2021), we generated predicted protein structures using these protein sequence datasets. AlphaFold2 generates quality metrics such as the predicted template modelling score (pTM; Jumper et al., 2021) that can then be used as a proxy for overall protein prediction confidence. We discarded structures with pTM values < 0.5 as these structures are less likely to be accurate.

### Filtering Tentative Allorecognition Proteins

For each of our cnidarian genomes of interest, we searched the predicted proteome sequences for likely homologs of the allorecognition protein family. We applied an initial filter for tentative homology by performing reciprocal BLASTP searches between our predicted proteome of interest and published allorecognition proteins from *H.* symbiolongicarpus (Huene et al., 2022). We restricted our tentative set proteins to those having bit-score values > 50 in reciprocal BLAST searches. We tested various cutoffs for bit-score and similar homology metrics (e.g., E-value) to assess tentative homology detection. Following optimization, this approach provided an appropriate cutoff for detecting homologs between closely related species. We further limited our candidates to those having a signal peptide, as detected by the SignalP webserver (Teufel et al., 2022). Proteins were also screened for likely structural homology using Foldseek (van Kempen et al., 2024) and DALI (Holm, 2022). Previous studies focused on remote homology detection emphasized the relative strengths of each methodology (Holm et al., 2023; van Kempen, 2024). Foldseek tends to run much faster but has a higher false positive rate, so combining both approaches should yield more accurate results (Himmel et al., 2023).

Using Foldseek, a set of predicted protein structures from each cnidarian of interest was compared to our database of published allorecognition domains (Huene et al., 2022) and the Protein Data Bank (PDB; Berman et al., 2000; Berman et al, 2003). Like AlphaFold2, Foldseek reports template matching (TM-score) quality metrics; here, we only considered high-quality matches, defined as those having a TM-score > 0.5.

Similarly, using DALI, we compared our cnidarian predicted proteome of interest to both our allorecognition domain dataset and a general protein dataset (AlphaFold2 in this case). DALI match results report root mean square deviation (RMSD) as a metric of dissimilarity between any two protein structures. Here, a cutoff of RMSD < 5 Å was used to define high-quality matches of interest, based on existing literature from protein docking simulations (Simons et al., 2001) and remote protein homolog comparisons (Kufareva & Abagyan, 2012). For synteny analyses, we used a more stringent RMSD < 2 Å cutoff to identify homologous relationships with higher confidence. Using these data, we limited our tentative allorecognition homologs to those with high-quality matches identified by both Foldseek and DALI, matching both the allorecognition domains and a general protein database (PDB for Foldseek, AlphaFold Protein Structure Database for DALI; Berman et al., 2000; Varadi et al., 2021).

### Structural Phylogenetic and Ortholog-Based Analyses

In parallel, we used two complementary approaches to assess evolutionary relationships among candidate ALRs: phylogenetic reconstruction and orthology-based clustering. Phylogenies were constructed using Foldtree (Moi et al., 2025), using our tentative filtered cnidarian ALRs, as well as with published *Hydractinia* ALRs (Huene et al., 2022), with trees constructed using default Foldtree parameters.

Orthogroup assignments in cnidarians were based on a previous OrthoFinder v2.5.4 run (Emms & Kelly, 2019), and we filtered these results for orthogroups in which at least one species had a tentative Alr gene; this allowed us to visualize orthogroups shared between cnidarian species as well as any species-specific orthologs. Using genome annotations available from NCBI, we visualized the positions of these genes in chromosomal space. Our chromosome homology assignments are based on previous macrosynteny analyses in cnidarians (Chang et al., 2026).

## Results

Our multi-pronged computational strategy, which combined both sequence-based homology detection (BLASTP) with structural comparison approaches (DALI and Foldseek), enabled the identification of candidate ALR homologs in multiple chromosome-length cnidarian genome assemblies that proved difficult to detect by sequence-based methods alone. This approach provided evidence of a *bona fide* ARC in three other cnidarian species: *H. vulgaris*, *P. americana,* and *N. vectensis*. In each species, genes encoding ALR homologs clustered within a contiguous genomic region syntenic to the *Hydractinia* allorecognition complex, supporting the existence of a conserved ARC across these cnidarian lineages.

Beyond identifying genomic clusters, this approach also resolved the domain architecture of candidate ALR homologs across cnidarian species. Multiple proteins identified in cnidarians such as *Podocoryna* had hallmark features associated with ALRs, including sets of tandem immunoglobulin-like domains (Fig 1). These proteins follow a pattern of domain organization similar to what is observed in *Hydractinia* ALRs, where a more variable V-set domain precedes a more invariant I-set domain (Huene et al., 2022). This arrangement is consistent with broader trends governing immunoglobulin-mediated protein-protein interactions and is similar to the pattern observed in immunoglobulin-like egg coat proteins, where the most N-terminal domains tend to be more variable (Luo et al., 2025; Rivera et al., 2022).

**Figure 1:**
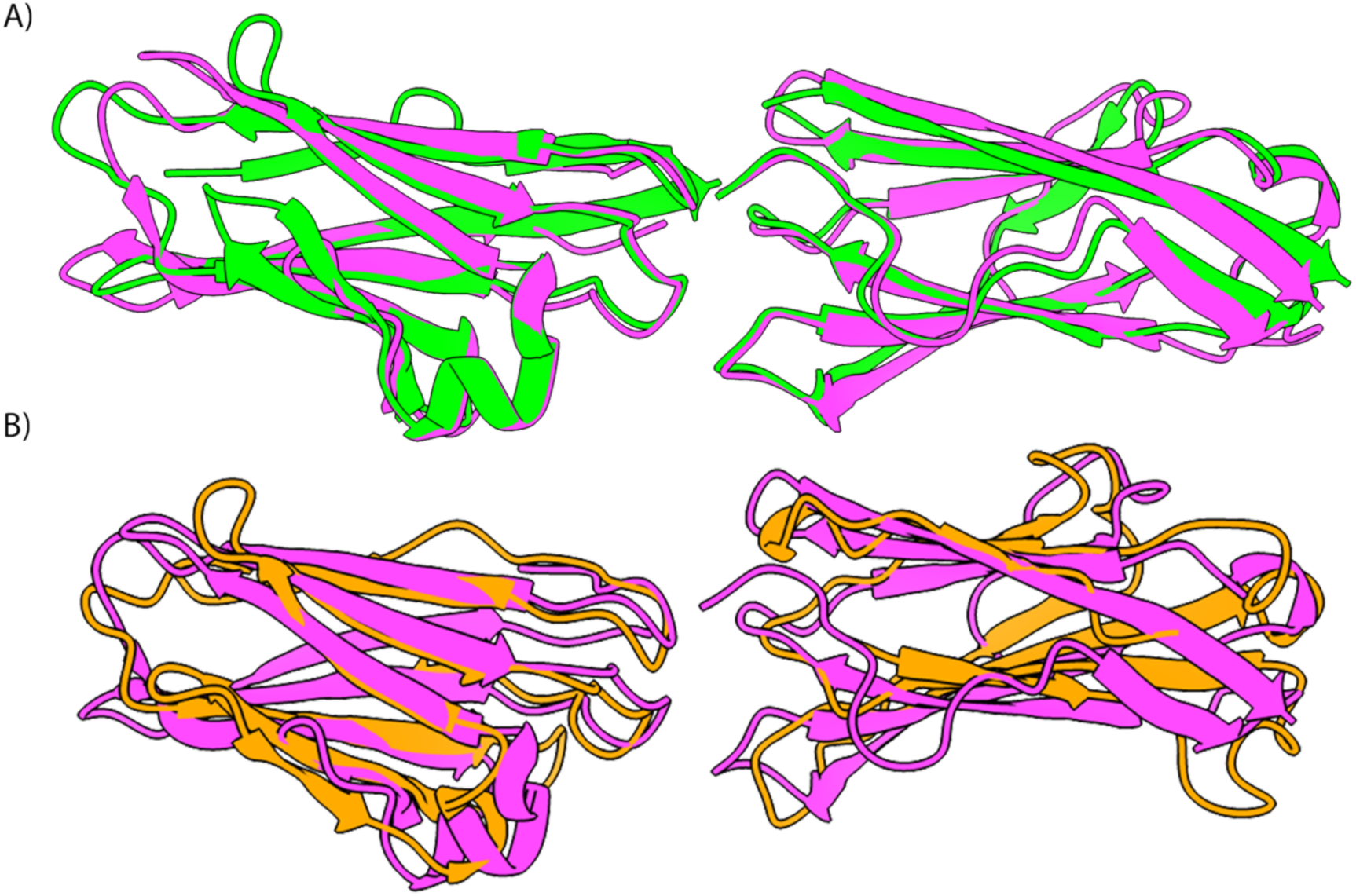
(A) Structural alignment of the published ALR1 structure from *Hydractinia symbiolongicarpus* (green) and a tentative *Podocoryna* ALR homolog (magenta). This alignment is comprised of the first two ALR domains, with a V-set domain (left) and an I-set domain (right). (B) Structural alignment between the same tentative *Podocoryna* ALR homolog (magenta) and a human heavy-chain only antibody crystal structure (orange, ID: 7KQY; Mieszkowski et al., 2020).

*Podocoryna* contains 22 tentative Alr genes, spanning approximately 18 Mb on chromosome 7 (Chang et al., 2026). This chromosome is homologous to *Hydractinia* chromosome 3 which contains its own allorecognition complex (Chang et al., 2026; Salamanca-Díaz et al., 2025). *Podocoryna* contains other allorecognition homologs outside of this gene complex, with four putative Alr genes on chromosome 4 and a single putative Alr gene on chromosomes 1, 6, 11, and 14 (Chang et al., 2026). While these genes do not fall within the gene complex, they may nonetheless represent *bona fide* allorecognition homologs. Tentative allorecognition homologs can display variation in the number of Ig-like domains they possess (Fig. 2), consistent with previous observations comparing *Hydractinia* ALRs (Huene et al., 2022).

**Figure 2:**
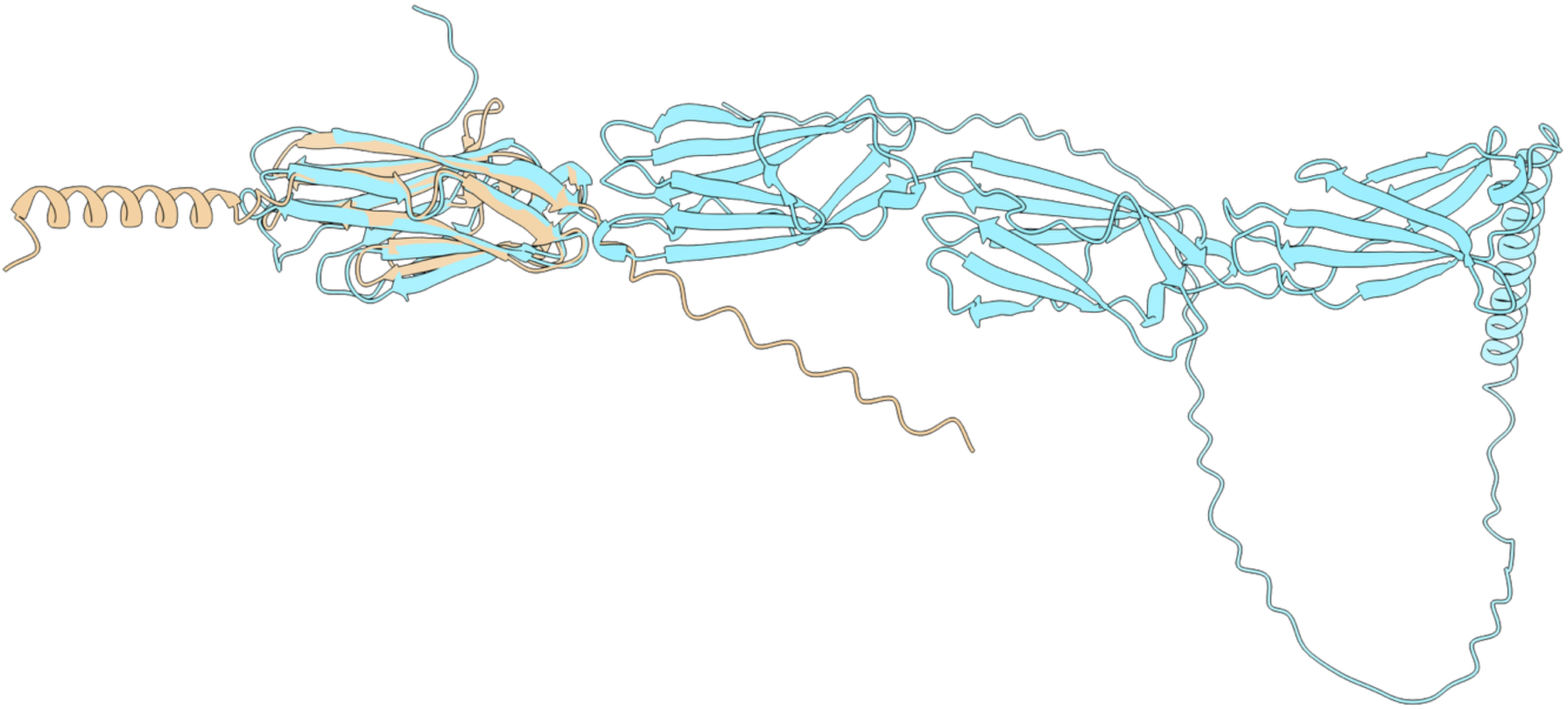
Predicted protein structures from two representative *Podocoryna* ALR homologs. These structures demonstrate variability in Ig-like domain number, consistent with the known *Hydractinia* Alr gene complex. Candidate proteins also differ in the N-terminal and C-terminal regions outside of these Ig-like domains. Blue, predicted homolog of Alr36; yellow, predicted homolog of Alr12A.

*Hydra vulgaris* also shows evidence of an allorecognition gene complex, with 10 tentative Alr genes on *Hydra* chromosome 1. This chromosome is also homologous to the chromosome containing the *Hydractinia* ARC. Genes encoding tentative homologs are also found outside of the predicted allorecognition gene complex, consistent with what is observed in both the *Podocoryna* and *Hydractinia* genomes (Chang et al., 2026). In addition, the *Hydra* ALR homologs are more dispersed, with four putative Alr genes on chromosome 13, three on both chromosomes 3 and 10, and a single predicted Alr gene on chromosomes 5, 6, and 14.

These tentative *Hydra, Podocoryna,* and *Nematostella* ARCs occur on chromosomes that are homologous to the chromosome containing the published *Hydractinia* allorecognition complex (Fig. 3), with this macrosynteny strongly suggesting that these gene complexes are truly homologous. Despite the presence of a conserved allorecognition complex, gene number and order do differ between hydrozoan species (Chang et al., 2026); this is not surprising given that gene collinearity is less conserved than chromosome-scale synteny (Schultz et al., 2023).

**Figure 3:**
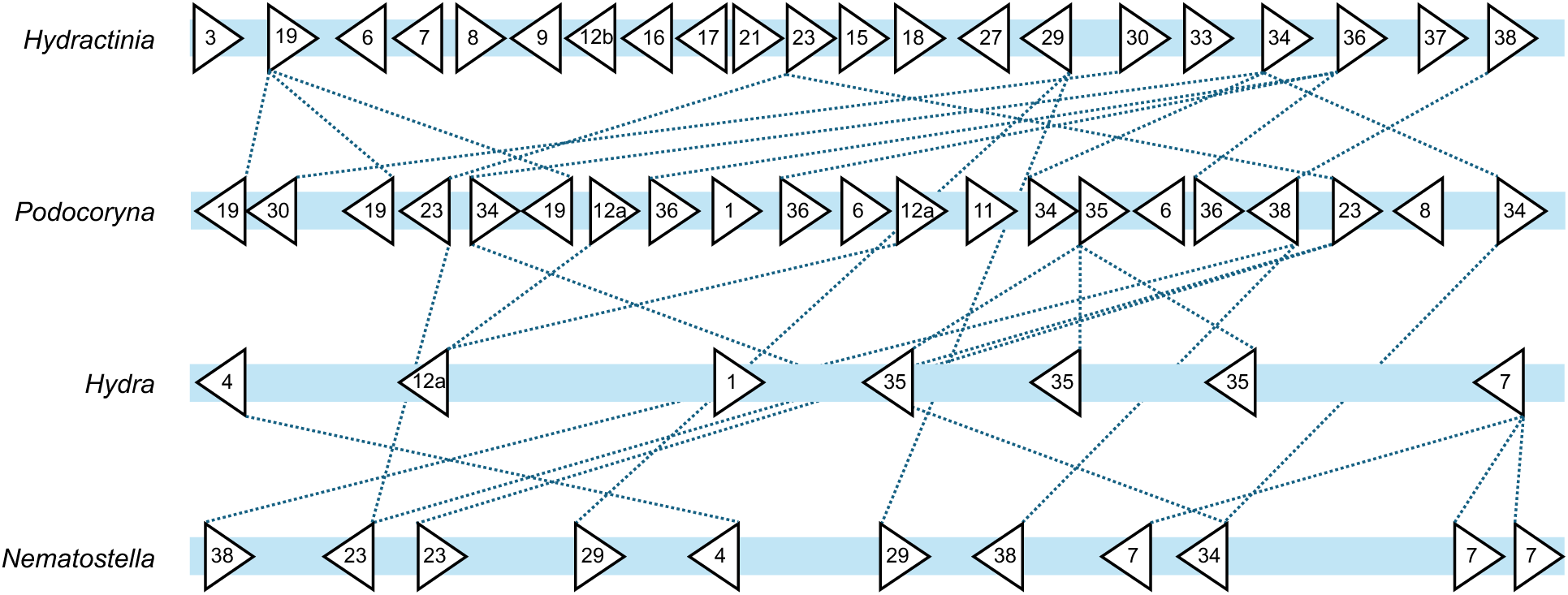
Predicted homologous Alr-like genes across homologous cnidarian chromosomes. We observe evidence of an allorecognition complex in *Hydractinia*, *Podocoryna*, *Hydra*, and *Nematostella*. However, local gene content and gene order can differ (Veltri et al., 2016). Numerical designations are used for protein names (e.g, ‘1’ for Alr1), and dotted lines connect predicted homologs between species.

Outside of the hydrozoans, we detected tentative ALR homologs in scyphozoans and anthozoan corals. In *Nematostella*, two chromosomes each contained 15 predicted Alr genes; one of these chromosomes is homologous to the chromosome containing the published *Hydractinia* ARC (Huene et al., 2022). Genes encoding an additional 26 putative ALR homologs are scattered across seven *Nematostella* chromosomes. Putative Alr genes were also detected in *Pocillopora damicornis* and *Rhopilema esculentum*; however, the NCBI-annotated versions of those genomes are not chromosome-scale, precluding further conclusions about gene complex organization in those species. While gene order and content can differ between the species, homology between the chromosomes points to a shared evolutionary origin.

It is possible that the genomic organization of Alr genes into an allorecognition gene complex is less conserved at greater taxonomic distances. Due to physiological differences, corals and free-living jellyfish may have a reduced need for this robust allorecognition function, or these genes may not need to be contiguous to perform their essential function. For instance, a free-living jellyfish can escape the fierce competition for space experienced by sessile hydrozoan colonies. Such differences in organismal lifecycles would affect the evolutionary pressures these species encounter and may have shaped the overall evolutionary history of the Alr gene family.

Our search for tentative ALR proteins in cnidarians identified multiple overlapping orthogroups across the species queried (Fig 4). Seven orthogroups are shared across all the taxa analyzed, which could suggest an ancestral set of allorecognition genes that is relatively conserved across all cnidarians. These shared orthogroups may represent those most essential for the allorecognition function and could be high-priority targets for further functional investigation. Lineage-specific gene loss may also have occurred, as comparative genomic analyses in eukaryotes suggest that major biological innovations may be associated with gene gain or loss at key moments in evolutionary history (Domazet-Lošo et al., 2024). Nine orthogroups are also shared between *Hydractinia* and *Podocoryna*, which could reflect an expansion of this gene family compared to their common ancestor. Finally, additional predicted orthogroups are observed in *Nematostella* than in its closest anthozoan relatives.

**Figure 4:**
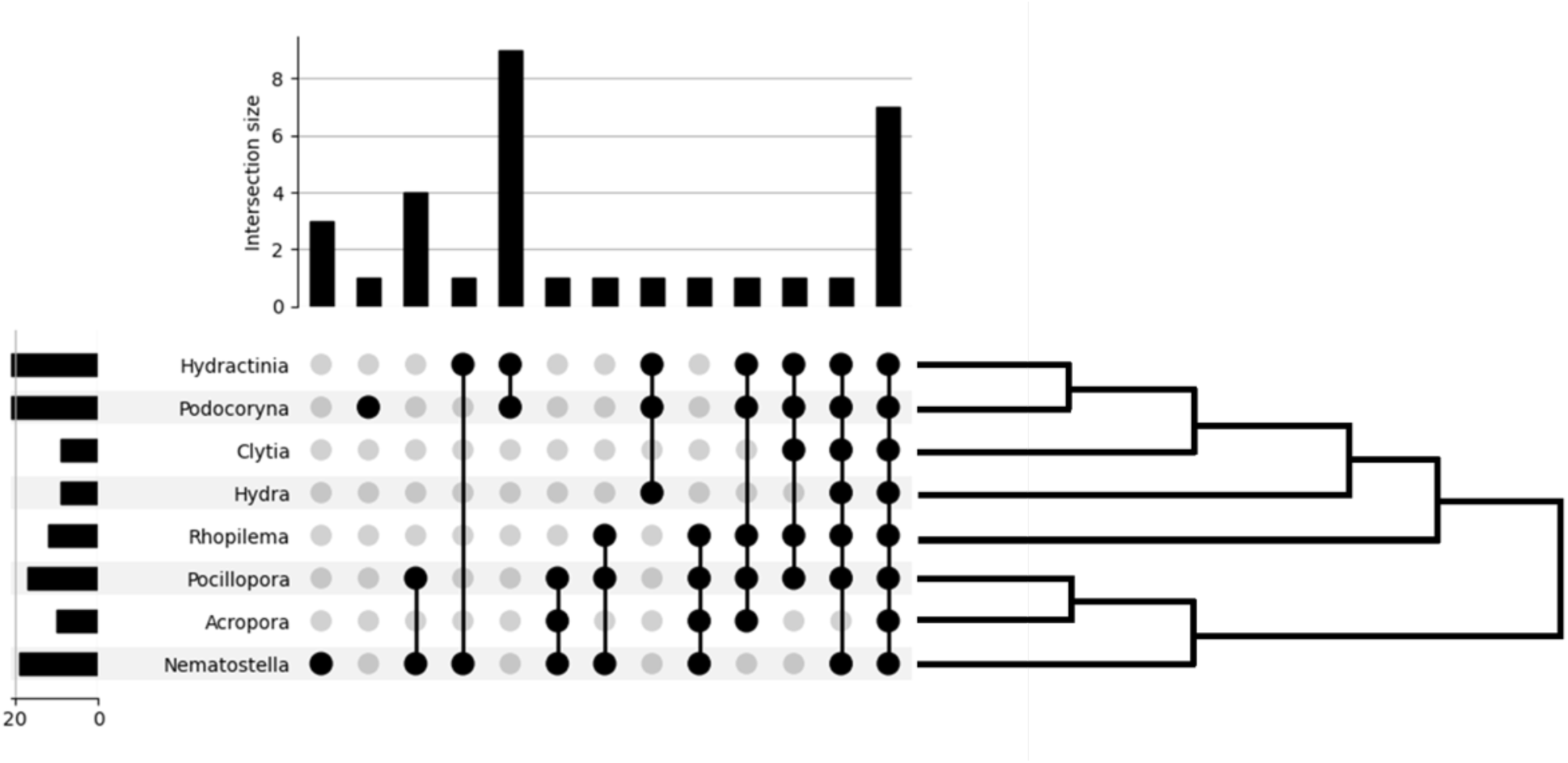
UpSet plot displaying the number of orthogroups shared by different cnidarian species. The scale to the left indicates the number of orthogroups present in each taxon, while the bar chart at the top indicates the number of orthogroups shared by multiple taxa. Seven orthogroups are shared across the eight cnidarians analyzed. There are also nine orthogroups that are found in both *Podocoryna* and *Hydractinia*.

Integrating our predictions with our OrthoFinder results (Emms & Kelly, 2019; Chang et al., 2026) allowed us to recover additional Alr-like genes not initially captured by our structure-based homology searches. Using this approach, we identified 20 tentative Alr genes in *Acropora millepora* that fell into 10 predicted orthogroups distributed across six chromosomes and two unplaced contigs in the *Acropora* genome.

The largest intersection of orthogroups is shared between *Hydractinia* and *Podocoryna*, which could reflect a lineage-specific gene expansion in the ancestor of these taxa. Gene loss elsewhere in the tree is also possible, though this would be a less parsimonious explanation for this observation. These findings raise important questions about the selective pressures governing Alr gene number across cnidarian species. Lifestyle differences may partly explain the variation observed: the stationary, colonial lifestyle of *Hydractinia*, which demands constant negotiation of space and resources with conspecifics, may drive selection for a larger and more diversified ALR repertoire, whereas the free-living medusa stage of *Podocoryna* (Miglietta & Cunningham, 2012) facilitates both sexual reproduction and long-distance dispersal, potentially relaxing the selective pressure to maintain a large allorecognition gene complex.

Phylogenetic analyses helped clarify the evolution of this gene family within hydrozoans and cnidarians more broadly (Fig 5). Given our emphasis on detecting distant homologs through structural similarity, we performed structure-based phylogenetics in Foldtree using individual protein domains (Moi et al., 2025). The inclusion of published *Hydractinia* ALR domains in the phylogeny allowed us to contextualize our findings within a framework of previously described domain structures. *Hydractinia* ALR domains define phylogenetic groupings of V-set Ig-like domains, I-set Ig-like domains, and fibronectin-like (FN3-like) domains, which may reflect ancestral ALR domain classes that have persisted throughout cnidarian evolutionary history. The V-set and I-set domains typically comprise the first two Ig-like domains in an ALR protein. Phylogenies of V-set and I-set domains show close grouping of *Hydractinia* and *Podocoryna* sequences, consistent with a lineage-specific expansion of this gene family in the hydrozoan ancestor. This expansion may explain why *Podocoryna* and *Hydractinia* have more predicted ALRs than observed in other clades (Fig. 4) and show macrosynteny between their gene complexes (Fig. 3).

**Figure 5:**
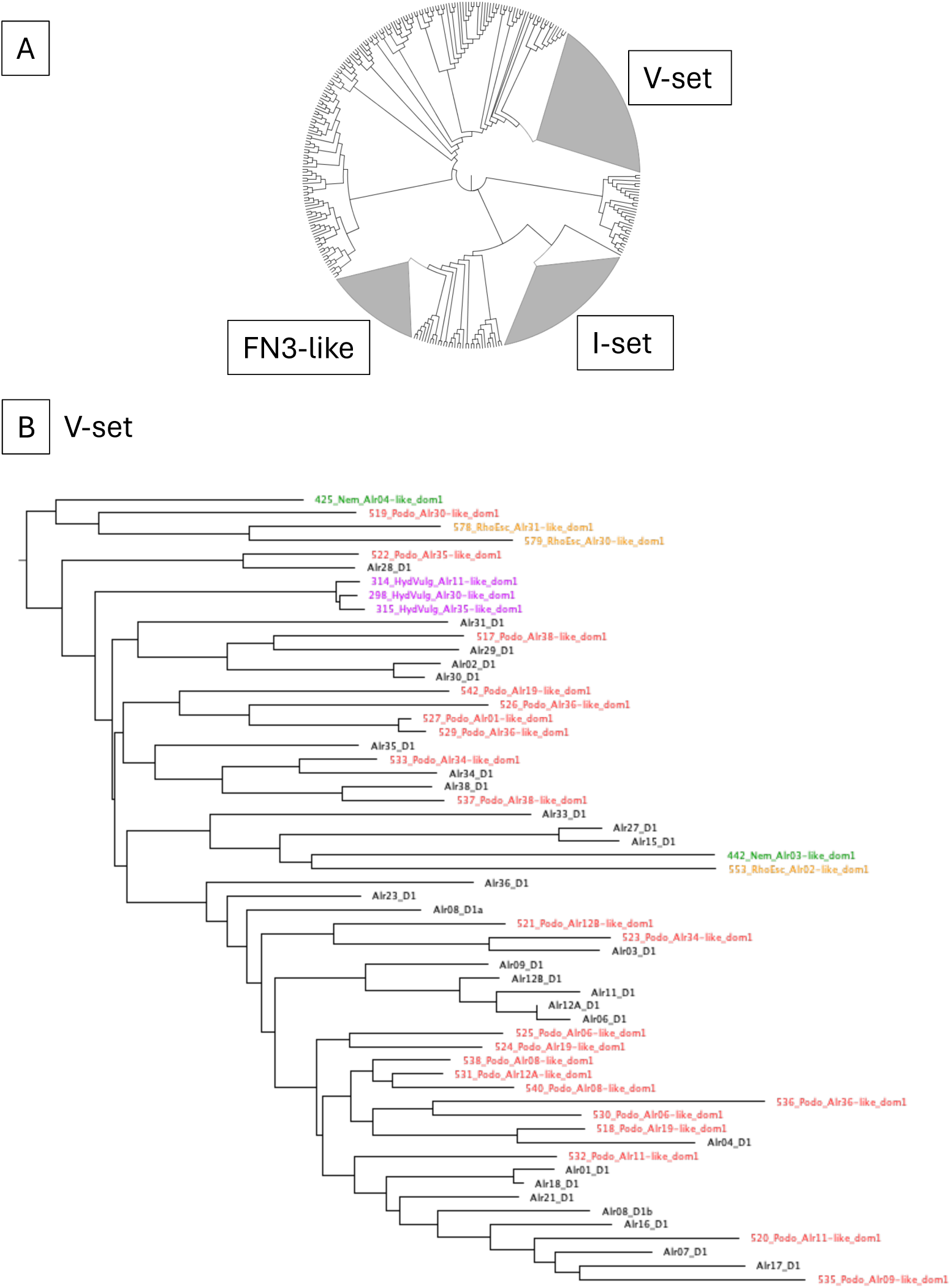
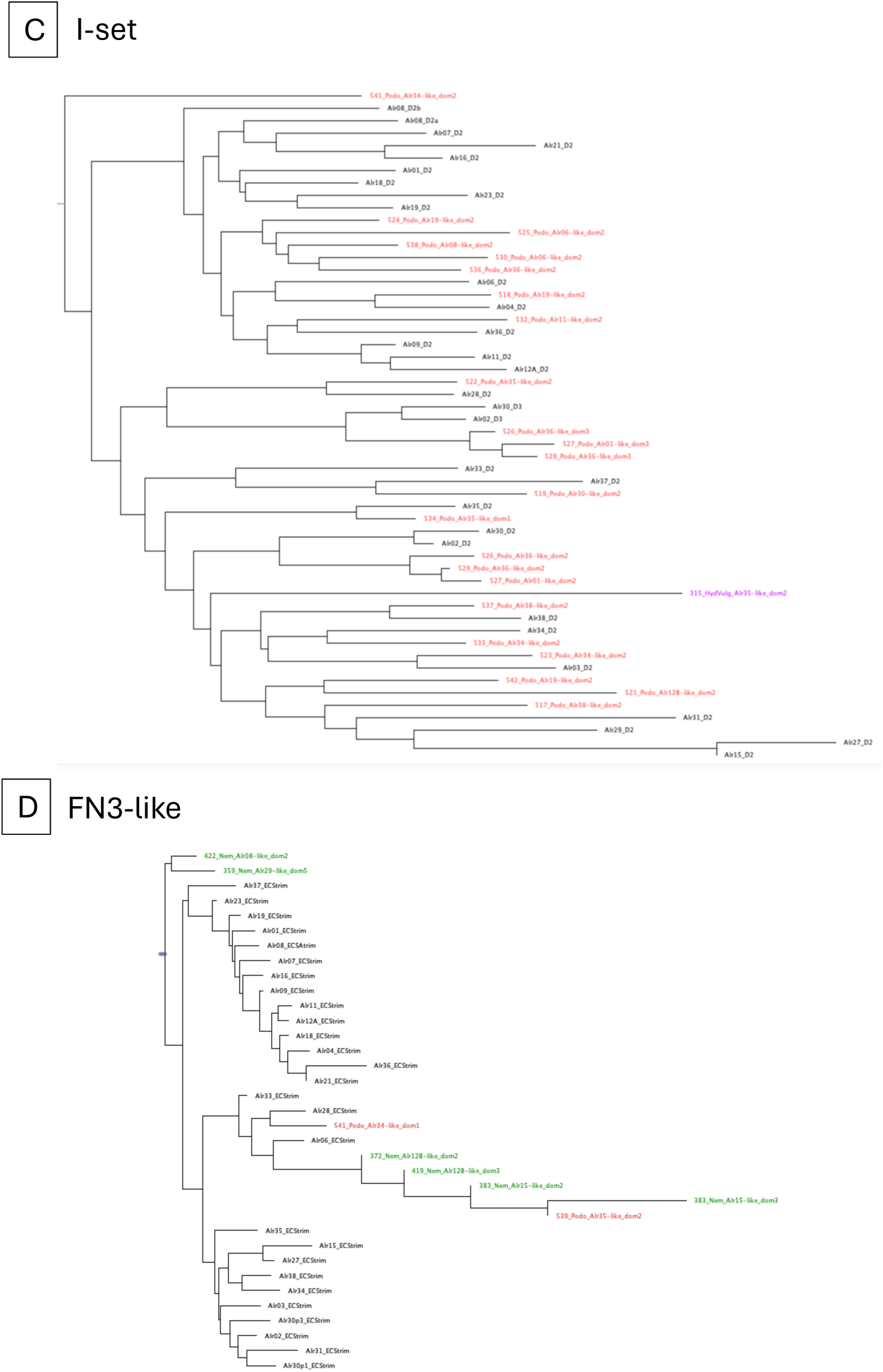
(A) Structure-based phylogeny constructed with Foldtree (Moi et al., 2025). V-set, I-set, and fibronectin-like (FN3-like) immunoglobulin domains form distinct phylogenetic clusters. (B, C) Phylogenies of the clades containing V-set and I-set domains, respectively. (D) Phylogeny of FN3-like domains. Black tip labels correspond to published *Hydractinia* ALRs. Tentative ALRs from *Podocoryna* (red), *Pseudorhiza* (brown), *Rhopilema* (orange), *Pocillopora* (blue), *Hydra* (violet), and *Nematostella* (green), are also displayed. We observe several clades with multiple species, as well as lineage specific gene expansions.

## Discussion and Conclusions

Collectively, our findings establish the allorecognition complex as a highly duplicated and deeply conserved gene family present across multiple cnidarian species. Although macrosynteny is maintained across vast evolutionary distances, gene collinearity and order can be lost relatively rapidly, a pattern clearly reflected in the evolutionary dynamics of the cnidarian allorecognition complex. While the complex consistently maps to homologous genomic regions in hydrozoans, gene number and order differ between species. Given its role in distinguishing self from non-self, this region may be subject to rapid evolutionary turnover, a dynamic observed in other systems where essential protein-protein interactions drive accelerated evolution, as has been shown for self/non-self recognition in amoebae (Holland et al., 2025). For instance, a mutation arising in an allorecognition gene could render a colony incompatible with an otherwise clonal neighbor, and such dynamics could facilitate the rapid reproductive isolation of new *Hydractinia* colonies and populations.

We further hypothesize that differences in life history strategy influence the evolution of this gene complex. As a strictly colonial and sessile organism, *H. symbiolongicarpus* likely faces strong selective pressure to maintain a robust allorecognition system (Huene et al., 2022; Nicotra, 2019; Rosengarten & Nicotra, 2011). In contrast, the closely related hydroid *Podocoryna* possesses a free-living medusa stage and fewer allorecognition genes; the medusa facilitates both sexual reproduction and long-distance dispersal, potentially relaxing selective pressure to maintain a large allorecognition gene complex. Whether the difference in Alr gene number between these taxa reflects substantial gene loss in *Podocoryna*, gene duplication in *Hydractinia*, or some combination of both remains an open question that will require further investigation into the ancestral state of the hydrozoan allorecognition complex. Consistent with this picture, our phylogenetic analyses indicate that certain ancestral ALR orthologs have been retained throughout cnidarian evolution. Furthermore, a lineage-specific gene expansion evident in both *Podocoryna* and *Hydractinia* is a likely contributing to the higher number of ALR orthologs observed in these species.

These findings also inform our broader investigation into the evolutionary relationship between invertebrate allorecognition and the bilaterian immune system. Although a clearly defined allorecognition gene complex is evident on homologous hydrozoan chromosomes, this pattern becomes less apparent at greater evolutionary distances, raising the possibility that not all bilaterian ALR homologs will map to homologous chromosomes. Syntenic regions remain the most tractable targets for remote homolog detection, though gene transposition to non-homologous chromosomes cannot be excluded. These considerations will guide future searches for remote homologs in the early-branching deuterostomes *Branchiostoma* and *Ciona* – closer relatives of vertebrates for which high-quality genome assemblies are available (Adkins et al., 2025; Rhie et al., 2021; Satou et al., 2019). Importantly, both species predate the two rounds of whole-genome duplication that occurred in the vertebrate stem lineage (Dehal & Boore, 2005; Huang et al., 2023), making them particularly informative outgroups for comparative homolog detection. The identification of *Branchiostoma* proteins with high structural similarity to hydrozoan ALRs lends support to this framework, and the integrated structural and genomic approach developed here is well positioned to address outstanding questions in the evolution of bilaterian immunity, including the deep origins of adaptive immune system genes in the MHC.

More broadly, the cnidarian ALR system represents yet another instance of a recurrent evolutionary pattern: the co-option of N-terminal V-set Ig-like domains to encode binding specificity in immune recognition. This pattern is evident across deeply divergent animal lineages, from the variable region-containing chitin-binding proteins (VCBPs) found in *Branchiostoma* (Cannon et al., 2002; Dishaw et al., 2011) to the fibrinogen-related proteins (FREPs) of lophotrochozoan invertebrates (Zhang & Loker, 2004) and possibly extending further still to the ancestral immunoglobulin domain architectures identified in sponges(Urdiciain et al., 2025). Taken together, these observations suggest that the co-option of Ig-like domains to generate immune recognition diversity is not a derived vertebrate strategy but instead reflects a deeply conserved solution that has shaped immune systems across the animal kingdom. Our approach demonstrates the potential of protein structure-based methods to elucidate the evolution of essential animal protein families.

## Notes

### Competing Interest Statement

The authors have declared no competing interest.

### Funding

- National Human Genome Research Institute, National Institutes of Health, Bethesda, MD US: ZIA HG000140

